# A genetic variation in the adenosine A2A receptor gene contributes to variability in oscillatory alpha power in wake and sleep EEG and A_1_ adenosine receptor availability in the human brain

**DOI:** 10.1101/2023.03.01.530170

**Authors:** Naemi L. Tichelman, Anna L. Foerges, Eva-Maria Elmenhorst, Denise Lange, Eva Hennecke, Diego M. Baur, Simone Beer, Tina Kroll, Bernd Neumaier, Andreas Bauer, Hans-Peter Landolt, Daniel Aeschbach, David Elmenhorst

## Abstract

The EEG alpha rhythm (8-13 Hz) is one of the most salient human brain activity rhythms. Spectral power in the alpha range in wakefulness and sleep varies among individuals based on genetical predisposition, yet knowledge about the underlying genes is scarce. The EEG alpha oscillations are related to cerebral energy metabolism and modulated by the level of attention and vigilance. The neuromodulator adenosine is directly linked to energy metabolism as product of adenosine tri-phosphate (ATP) breakdown and acts as a sleep promoting molecule by activitating A_1_ and A_2A_ adenosine receptors. We quantified EEG oscillatory alpha power in wakefulness and sleep, as well as A_1_ adenosine receptor availability by positron emission tomography with ^18^F-CPFPX, in a large sample of healthy volunteers carrying different alleles of gene variant rs5751876 of *ADORA2A* encoding A_2A_ adenosine receptors. Oscillatory alpha power was higher in homozygous C-allele carriers (n = 27, 11 females) compared to heterozygous and homozygous carriers of the T-allele (n(C/T) = 23, n(T/T) = 5, 13 females) (F_(18,37)_ = 2.35, p = 0.014, Wilk’s Λ = 0.467). Across considered brain regions an effect of *ADORA2A* genotype on A_1_ adenosine receptor binding potential was found (F_(18,40)_ = 2.62, p = 0.006, Wilk’s Λ = 0.459) and after correction for multiple testing this effect was shown to be significant for circumscribed occipital region of calcarine fissures.

A correlation between individual differences in oscillatory alpha power and adenosine receptor availability was found for the subgroup of female participants only. In conclusion: a genetic variation in the adenosinergic system affects individual alpha power, although a direct modulatory effect via the A_1_AR has been demonstrated for females only.

## 1. Introduction

Electroencephalography (EEG) captures brain activity in real-time and is frequently used in clinical diagnostics and research. Derived potential fluctuations are correlates of synchronized firing of cortical neuronal masses (Niedermeyer, 1996; Nunez & Srinivasan, 2006) and rhythmic oscillations are characterized by their frequency range, amplitude, and power (Proekt, 2018). Alpha waves, first described by Hans Berger in the 1920s are (since then) intensively studied with constantly evolving techniques (Bazanova & Vernon, 2014). Ranging from 8 - 13 Hz, alpha oscillations are recorded predominantly in the state of rested wakefulness, most evidently at parieto-occipital derivations (Brown et al., 2012). As a basis of cognitive brain functions like attention (Saalmann et al., 2012), perception (Busch et al., 2009; Samaha & Postle, 2015), and working memory (Klimesch, 1996), alpha activity possibly enables effective neuronal processing by inhibiting uninvolved brain areas (Jensen & Mazaheri, 2010; Klimesch, 2012; Sadaghiani & Kleinschmidt, 2016). With increasing arousal levels or attentional demand, alpha waves are attenuated and replaced by faster EEG rhythms in the beta and gamma range (Nunez & Srinivasan, 2006). At the transmission to sleep, alpha waves diminish while during sleep slower oscillations and specific sleep patterns dominate the EEG (Brown et al., 2012). When looking at rhythmic oscillations in neural powerspectra it is important to differentiate between the aperiodic and periodic component, which are two distinct functional processes (Donoghue et al., 2020). For instance, Ouyang and colleages demonstrated that after separating arrhythmic activity reflecting 1/f-like characteristics and alpha oscillations, the aperiodic parameters, not alpha power, predicts between-individuals variability in cognitive speed (Ouyang et al., 2020).

Several studies, in cell cultures and animals, suggest that alpha oscillations are generated mainly from intracortical pyramidal neurons (Lopes da Silva, 1991). Propagated by an interplay of relay neurons in the thalamus (Hughes & Crunelli, 2005; Lopes da Silva, 1991; Saalmann et al., 2012) and cortico-cortical feedback-loops, alpha activity is modulated by intracortical and thalamo-cortical transmission as well as by cholinergic input from the brainstem (Hughes et al., 2004). Vigilance, as well as homeostatic and circadian sleep-wake regulatory processes, impact brain oscillations such as the alpha rhythm and should therefore be considered when investigating their underlying nature. Indeed, EEG-frequency-dependent homeostatic and circadian components have been extracted from power spectra during extended wakefulness, and the kinetics of the wake-dependent increase of theta-low-frequency alpha power (5-9 Hz) was found to reflect the changes of the homeostatic process previously derived from the sleep EEG (Aeschbach et al., 1999; Aeschbach et al., 2001).

Adenosine, a neuromodulator and component of cellular energy metabolism, has various physiological functions including an important role in sleep homeostasis (Bjorness et al., 2009; Thakkar et al., 2008). According to the two-process model, a homeostatic process regulates sleep onset and maintenance through interaction with an underlying circadian process (Borbely, 1982). As a sleep factor, whose concentration increases as a function of duration of wakefulness, adenosine initiates and preserves sleep (Porkka-Heiskanen et al., 1997; Strecker et al., 2000). It modulates neuronal transmission by four different subtypes of G protein-coupled receptors, whose subtypes A_1_AR and A_2A_AR are most widely distributed in the brain (Ribeiro et al., 2002). Cellular mechanisms of adenosine signaling include presynaptic inhibition of cortical and thalamic neurons (Pape, 1992), reduction of the cholinergic input from the brainstem (Porkka-Heiskanen et al., 1997; Rainnie et al., 1994) and inhibition of wake-promoting neurons in the basal forebrain (Lazarus et al., 2019; Lazarus et al., 2022).

Genetic variation of the gene encoding the A_2A_AR (*ADORA2A*, c.1976T>C polymorphism, rs5751876) was shown to be associated with inter-individual differences in anxiety response to caffeine (Alsene et al., 2003), sensitivity to wake-promoting effects of caffeine (Bodenmann et al., 2012; Retey et al., 2007) and cognitive performance after sleep deprivation (Bodenmann et al., 2012; Urry & Landolt, 2015). This allele variant in the *ADORA2A* gene was found to have an impact on cerebral A_1_AR availability (Hohoff et al., 2014; Hohoff et al., 2020). Alpha activity during rested wakefulness and sleep varied significantly between different *ADORA2A* allele carriers (Retey et al., 2005).

In summary, adenosine signaling is involved in the regulation of sleep-wake transition and its receptors seem to be involved in neuronal activity both during wakefulness and sleep. EEG activity including alpha power varies considerably among individuals and is genetically influenced in wakefulness and sleep (Ambrosius et al., 2008; De Gennaro et al., 2008; Salmela et al., 2016; van Beijsterveldt & van Baal, 2002). Current clinical research aims at the identification of non-invasive and affordable biomarker tools such as the EEG, with a focus on the diagnosis of neuropathologic states and neurological and psychiatric diseases (Cassani et al., 2018).

Here, we investigate aspects of inter-individual variability in EEG characteristics, in particular alpha power, in order to improve our understanding of EEG features that might have diagnostic potentials and open new therapeutic possibilities. We determined A_1_AR availability in a sample of healthy subjects and recorded EEG at rested conditions. Group differences between allele carriers of the *ADORA2A* rs5751876 variant were analyzed. As adenosine signaling modulates neuronal transmission, the availability of A_1_ARs in cortical brain regions could have a direct impact on electrophysiological measures at near-by derivations. A distant effect of adenosine signaling on alpha activity at scalp level could be triggered by an attenuating effects on alpha-generating neurons in the thalamus.

Therefore we aimed at characterizing the modulatory effect of gene variant rs5751876 of *ADORA2A* by specifically addressing the following questions: (1) Does this variant modulate features of the EEG in the conditions of restful waking and sleep, in particular oscillatory alpha power? (2) Does this variant account for inter-individual differences in the binding potentials of the A_1_AR? (3) Do inter-individual differences in A_1_ARs availability in cortical and subcortical brain regions correlate with inter-individual disparities in oscillatory alpha power?

## 2. Methods

All experimental procedures were approved by the Ethics committee of the Medical Association of North Rhine, Germany, and the German Federal Office for Radiation Protection. They were conducted according to the principles of the Declaration of Helsinki. All study participants gave written informed consent.

### 2.1 Participants and genetic testing

EEG and PET data of 59 healthy volunteers (26 females; range 20 – 39 years; mean age 27 ± 5 years) were included in the analysis. The data presented here were from two different studies. The first study (study #1) aimed to investigate chronic sleep deprivation and its effects on cognitive performance, attentional abilities and cerebral A_1_AR availability. The second study (study #2) with a similar design focused additionally on exploring effects of caffeine combating fatigue and performance decrements in a selected sample of healthy, homozygous C-allele carriers of the *ADORA2A* rs5751876 gene variant. Respective methodological details have been published for study #1 (Hennecke et al., 2021) and study #2 (Baur et al., 2021). Recruited volunteers reported a habitual sleep duration between 6 - 9 h, no sleep-wake disorder or chronic disease, no history of head injury, substance abuse, or intake of any medication (except contraceptives). Participants were non-smokers, right-handed and reported estimated average daily caffeine intake of no more than 450 mg. For determination of the c.1976T>C polymorphism (SNP-ID: rs5751876) blood (study #1) or salvia samples (study #2) were analyzed. Genomic DNA was extracted from samples and submitted to the genotyping procedure as previously described (Baur et al., 2021). Briefly, allele-specific primers were applied to amplify each allele of concern during the PCR procedure and allelic identification was performed automatically by SDS software (version 2.2) of the Applied Biosystems PRISM 7900HT.

### 2.2 Study protocols

Experimental procedures at consecutive days and nights took place in the sleep laboratory of the German Aerospace Center, Cologne (https://www.dlr.de/envihab/). At least one week before admission participants were asked to maintain a regular sleep schedule (time in bed (TIB) from 10:00 PM to 7:00 AM or 11:00 PM to 8:00 AM (study #1) and 11:00 PM to 7:00 AM or 12:00 AM to 8:00 AM (study #2), no daytime naps) and abstain from caffeine and alcohol. To check compliance, subjects wore an actigraph on their non-dominant wrist during all time. All participants completed one adaption night and two baseline nights (8 h TIB). In both study protocols, daily assessment of cognitive performance, waking EEG recording, and polysomnography were conducted.

The sleep EEG data of 8-h TIB sleep period as well as wake EEG and PET data obtained on the next day were included in the present analysis.

### 2.3 Waking and sleep EEG recording

Waking resting-state EEG recording was performed twice a day. A high-density device (256-electrode-HydroCel Cap with an MRI-compatible Net Amps 300 amplifier; Electrical Geodesic Inc., Eugene, OR) was used for data acquisition. For 5-min-records impedances were kept below 100 kΩ. EEG signals were digitized at a sampling rate of 250 Hz, referenced to the vertex electrode (Cz). Recording took place in a quiet examination room with dimmed light (<100 lx). Subjects were instructed to sit in an upright comfortable position, to not move their head and muscles, and to fixate a black dot on the white wall in front of them during data collection. They were also instructed to avoid blinks, or otherwise blink in clusters in order to minimize EEG artifacts. To minimize distraction, the investigator left the room after starting the recording.

Subjects underwent continuous polysomnography during all nights in the laboratory. EEG signals at 6 bipolar derivations (according to the recommendations of the American Academy of Sleep Medicine (AASM): F4-A1, C4-A1, O2-A1, F3-A2, C3-A2, O1-A2) at a sampling rate of 256 Hz as well as electrocardiography, electromyography, finger pulse and breathing were measured. EEG signals were amplified with a time constant of 2.3 s and low-pass filtered with −6 dB at 70 Hz (Hennecke et al., 2021).

### 2.4 Waking and sleep EEG signal processing and spectral analysis

Wake EEG data were further processed off-line using Net Station 5 software (Geodesic EEG Software, version 5.4.2). Channels with consistent artifacts were marked manually and interpolated from the surrounding channels with an integrated tool, which uses spherical splines (Net Station tools, Bad Channel Replacement). Data were re-referenced to a common average (of all channels) and band-pass filtered (0.1-40 Hz) (Proekt, 2018). To control for EEG contamination (due to eye movement artifacts etc.), signals were epoched and inspected visually in 8 representative channels (F3, F4, C3, C4, O1, O2, two EOG-channels) using an in-house LabView program (LabView, version 16.0f5). Selected 2-s epochs with blinks or other artefacts like muscle twitching were deleted from all channels. On average 92.20 ± 30.47 s (range 33-202 s) artifact-free EEG data per record were further analyzed. If fewer than 30 epochs were artifact-free, this record was excluded from further analysis (Babiloni et al., 2020). For spectral analysis, cleaned EEG data were subjected to Fourier transform (FFT) using a Hanning window of 2-s (no overlap), resulting in a resolution of the power spectrum of 0.5 Hz and a range of 0.5 - 25.0 Hz for each derivation. Power estimates were averaged across epochs and across defined electrode clusters corresponding to relevant sites of the 10/20 Electrode Placement System (Table 1). An average of the two records on the same day was considered to compensate for marginal power fluctuations related to changes in attentiveness.

**Table 1.** Electrodes of the HydroCel Net clustered according to their 10-20 position and considered nearby brain regions

| **<u>Regional power values</u>** |  | **<u>Cortical brain regions that are located close to derivations</u>** |
| --- | --- | --- |
| **<u>Clustered electrodes # of the high-density net</u>** |  |  |
| F3 | 36, 40, 41, 42, 49, 50 | Sup. frontal gyrus , Mid. frontal gyrus, Inf. frontal gyrus |
| F4 | 224, 223, 214, 206, 213, 205 | Sup. frontal gyrus, Mid. frontal gyrus, Inf. frontal gyrus |
| F7 | 39, 46, 47, 48, 54 | Rolandic operculum, Mid. frontal gyrus |
| F8 | 3, 10, 2, 222, 1 | Rolandic operculum, Mid. frontal gyrus |
| Fz | 21, 22, 14 | Sup. frontal gyrus, Gyrus rectus |
| C3 | 51, 52, 58, 59, 60, 65, 66 | Precentral, Paracentral lobule |
| C4 | 196, 184, 195, 183, 155, 182, 164 | Precentral, Paracentral lobule |
| Cz | 9, 186, 132, 81, 8, 17, 44, 53, 80, 90, 131, 144, 185, 198 | Precentral gyrus, Postcentral gyrus, Paracentral lobule |
| P3 | 76, 77, 85, 86, 87, 97, 98 | Postcentral gyrus |
| P4 | 172, 163, 171, 162, 153, 161, 152 | Postcentral gyrus |
| P5 | 84, 94, 95, 96, 105 | Supramarginal gyrus, Angular gyrus |
| P6 | 179, 190, 178, 170, 177 | Supramarginal gyrus, Angular gyrus |
| Pz | 101, 129, 128, 119, 110, 100 | Precuneus |
| O1 | 108, 107, 117, 116, 115, 123, 124, 136 | Cuneus, Occipital lobe, Lingual gyrus |
| O2 | 150, 139, 151, 160, 159, 158, 149, 148 | Cuneus, Occipital lobe, Lingual gyrus |
| Oz | 126, 118, 127, 137, 125, 138 | Calcarine fissure |
| T7 | 63, 68, 69, 70, 74 | Temporal lobe |
| T8 | 203, 210, 202, 193, 192 | Temporal lobe |

Based on the polysomnographic recordings, sleep stages were scored in 30 s epochs according to the criteria of the American Academy of Sleep Medicine (Berry et al., 2017). After manually removing epochs with movement-artifacts, power spectra were computed with an FFT routine using a 4-s Hanning window and averaging ten overlapping epochs into 30-s epochs that matched the sleep scores (Elmenhorst et al., 2022). Power values were averaged across all epochs of each sleep stage, with non-rapid eye movement (NREM) stage comprising all N2 and N3 scored epochs. Power values of derivation pairs over both hemispheres (C3-A2 and C4-A1, etc.) were averaged. Absolute alpha power density was calculated by averaging power estimates in the range of 8-13 Hz for waking EEG and in the range of 8-11.5 Hz for sleep EEG.

Next to absolute spectral estimates we aimed to extract oscillatory alpha power correcting for the 1/f-like aperiodic component. Therefore the FOOOF algorithm (fitting oscillations & one over f, version 1.0.0) was used to parameterize neural power spectra. Settings for the algorithm were set as: peak width limits: [1, 6]; max number of peaks: 5; minimum peak height: 0.0; peak threshold: 2; and aperiodic mode: ‘fixed’ (Donoghue et al., 2020). Oscillatory alpha power was defined as peak power for the individually found center frequency in the alpha range. For wake EEG alpha range was defined from 8 to 13 Hz, while for sleep EEG the range 8 to 11.5 Hz was chosen to not be confounded by spindle activity which typically lays in the range of 12 to 13 Hz. Based on the metrics for goodness of the fit, participants with an error > 0.17 were excluded.

### 2.5 [^18^F]CPFPX PET data acquisition and processing

A_1_AR availability was quantified using [^18^F] CPFPX positron emission tomography (PET) as described previously (Elmenhorst et al., 2017; Elmenhorst et al., 2018).

PET and high-resolution three-dimensional T_1_-weighted magnetic resonance (MR) imaging were performed on an integrated 3 Tesla whole-body PET/MR system (Biograph mMR, Siemens Healthineers) (Delso et al., 2011) at the:envihab laboratory of German Aerospace Centre. Daily calibration and normalization of the PET scanner were performed using a ^68^Ge/^68^Ga phantom and cross-calibration factor was determined with a γ-counter (Wizard^2^; PerkinElmer). Synthesis and formulation of the radiotracer were done as described in (Holschbach et al., 2002) achieving a chemical purity of over 96 %. Diluted in sterile saline solution (0.9 %), the tracer was intravenously injected with an initial bolus (15.9 ml in 2 min) followed by constant infusion (34.1 ml in 118 min) with a Kbol value of 55 min (Elmenhorst et al., 2007). Scanning and injection were started simultaneously and scan duration was 100 min. Mean injected dose of [^18^F]CPFPX was 179.44 ± 20.782 MBq (range 103 - 200 MBq) with mean molar activity of 110.54 ± 73.651 GBq/μmol (range 21.56 - 317.62 GBq/μmol) and a corresponding average mass of injected CPFPX of 2.40 ± 1.55 nmol (range 0.37 - 8.53 nmol). For PET signal reconstruction the e7 tool (Siemens Molecular Imaging) was used, which applied the OP-OSEM reconstruction algorithm with point spread function modelling with 3 iterations and 21 subsets. For post-filtering, a 3 mm Gaussian filter was used and the framing scheme was 4 x 60 s, 3 x 120 s, 18 x 300 s. The resulting matrix dimensions were 344 x 344 x 127 with a reconstructed image resolution of 2.09 x 2.09 x 2.03 mm^3^. Images were corrected for detector normalization, randoms and scatter. A template-based attenuation correction was done based on the method described in (Izquierdo-Garcia et al., 2014).

Pre-processing of PET and corresponding MRI data was performed using the PMOD Neuro Tool (version 4.006; PMOD Technologies) as described previously (Pierling et al., 2021). Motion correction was applied by creating a reference image from average PET data of the first 9 min of the scan. Default matching parameters included squared difference sum cost function, trilinear interpolation and smoothing with 6 mm full width at half maximum (FWHM). When automatic segmentation was unsuccessful, MR images were cropped with an automatic function which removed the neck and restricted the MR data set to skull and brain. To segment T_1_-weighted MR images into grey matter (GM), white matter (WM), and cerebrospinal fluid, the images were denoised at medium strength and the 6 Probability Maps (SPM12) variant was applied. Further sampling parameter was set to 3.0 mm, bias regulation balanced for light modulations of the image intensity across the field-of-view and variations were smoothed using 60 mm FWHM. The clean-up was set at thorough and affine regularization was put according to European brains. For segmentation touch-up a background 0.2 probability level and overlay strategy with thresholds for GM and WM probability map of 0.1 and 0.05 was applied. When PET-MR matching was required, a rigid matching using normalized mutual information criterion with matching sampling of 3.0 pixel was done. For spatial normalization, probability maps transformation, informed by normalization results from previous MR segmentation, was used. The automated anatomical labelling template of the Montreal Neurological Institute which is implemented in the PMOD software (Tzourio-Mazoyer et al., 2002) defined volumes of interest (VOI). To check the borders of cortical VOIs and avoid false signal detection from cerebral sinuses, PET images in atlas space and GM probability information from the segmentation resulting mask were rated and manually adjusted if needed. In addition, cerebellar VOIs were manually adapted to ensure its usage as reference region, defined by a low A_1_AR availability (Bauer et al., 2003; Fastbom et al., 1987; Meyer et al., 2007).

Regional time-activity curves (TACs) were calculated for each VOI with the PMOD Kinetics Tool (version 4.006; PMOD Technologies). BP_ND_ was calculated according to Logan’s reference tissue model (t* = 30 min; (Logan et al., 1996)) using the cerebellum as reference region and an average k2’, resulting from the simplified reference tissue model.

### 2.6 Statistical analysis

SAS software (SAS Institute, Cary, NC) was used for statistical testing. To ensure validity and reliability of waking EEG records and processing steps, individual power spectra of every participant were critically inspected, absolute alpha power density values of the two time points (morning and afternoon) were checked for inter-correlation. Normal distribution of A_1_AR binding potentials was confirmed using the Shapiro-Wilk-test. For comparison study data were divided into two groups based on genotype (*ADORA2A* rs5751876): homozygous C-allele carriers (n = 31, 13 females) and homozygous or heterozygous carriers of the T-allele (n = 27, 13 females, 5 T/T-allele carriers). Oscillatory alpha power were subjected to a multivariate analysis of variance (MANOVA) with genotype-group as independent and oscaillatory alpha power at different scalp sites as dependent variables. Sleep EEG oscillatory alpha power at frontal, central and occipital scalp sites for NREM and REM sleep stages was subjected to separate MANOVA. The same analyses were done for absolute alpha power density values of wake and sleep EEG. Binding potentials of 18 atlas-defined brain regions were analyzed for an effect of *ADORA2A* rs5751876 allele variant applying MANOVA. P-values for oscillatory alpha power and A_1_AR binding potentials were corrected for multiple testing using Benjamini-Hochberg procedure. Local power values were organized in groups for topographical analysis in respect to nearby cortical brain regions (see Table 1). Explorative Pearson correlational analyses of A_1_AR BP_ND_ and oscillatory alpha power was performed for cortical VOIs which indicated variation for different *ADORA2A* variant carriers and nearby electrodes. A distant effect of A_1_AR amount on scalp EEG parameters was evaluated by correlation of A_1_AR availability in subcortical brain region of thalamus with oscillatory alpha power at central and occipital derivations. We tested for correlation in the whole sample (n = 59) as well as in subgroups of females and males.

## 3. Results

Demographic details of the participants are listed in Table 2. The two genotype-based groups did not differ in sex or age distribution, body mass index (BMI), injected tracer mass or activity and total sleep time (assessed by polysomnography).

**Table 2.** Demographic data and PET characteristics of C/C carriers versus C/T and T/T carriers: no significant differences between the groups concerning sex, age, or BMI. Activity of injected tracer-substance und sleeping period preceding the scan did not differ.

| <u>Demographics/characteristics (mean ± STD)</u> | <u>C/C-carrier, n= 31</u> | <u>C/T- & T/T-carrier, n=28</u> | <u>p value</u> |
| --- | --- | --- | --- |
| Sex (male/female) | 18/13 | 15/ 13 |  |
| Age (years) | 27 ± 5 | 27 ± 5 | 0.687 |
| Body mass index (BMI) | 23.56 ± 2.66 | 22.73 ± 2.07 | 0.191 |
| Injected activity (MBq) | 175.2 ± 21.99 | 184.5 ± 18.17 | 0.083 |
| Amount of substance (nmol) | 2.49 ± 1.76 | 2.26 ± 1.29 | 0.564 |
| Specific activity at scantime (GBq/μmol) | 108.4 ± 77.41 | 115.1 ± 70.05 | 0.733 |
| Time in bed (min) | 479.9 ± 1.34 | 480.3 ± 0.68 | 0.164 |
| Total sleep time (min) | 423.0 ± 32.09 | 421.9 ± 33.24 | 0.892 |

### 3.1 Absolute and oscillatory alpha power in sleep and wake EEG

We found ranging absolute alpha power densities depending on electrode position in well-rested conditions with highest powers measured for parieto-occipital leads. Intra-individual stability of absolute alpha power density could be demonstrated, as the values of the morning and afternoon records were highly correlated (for each scalp site p < 0.001, r = 0.703 – 0.925, n = 54 due to 5 subjects missing a second record). Inter-individual variability of EEG activity was indicated by the broad range for alpha power density levels in the sample.

Concerning oscillatory and aperiodic parameters extracted with the FOOOF tool, waking EEG data of three participants was excluded due to high error-values for the fit (> 0.17). Comparing median values, during sleep and waking, subjects indicated highest oscillatory alpha activity during waking (0.911μV^2^ at Oz scalp site, n=56) followed by rapid eye movement (REM) sleep, and NREM sleep (REM sleep 0.530μV^2^ vs. NREM sleep 0.255μV^2^, at occipital scalp site, n=59). With progress to deeper sleep stages (N1 to N3), oscillatory power in the alpha range decreased.

### 3.2 Alpha activity levels in C/C-allele carriers versus C/T- and T/T-allele carriers

In the waking EEG, MANOVA demonstrated a statistically significant global effect of *ADORA2A* genotype (F_(18,37)_ = 2.35, p = 0.014, Wilk’s Λ = 0.467) on oscillatory alpha power across 18 scalp sites (see Figure 2). Post hoc comparisons identified this effect to be significant for 13 of 18 considered scalp sites (p < 0.036, Benjamini-Hochberg corrected significance level). C/C-allele carriers had higher oscillatory alpha power compared to C/T- and T/T-allele carriers, especially at central, parietal, and occipital leads. For homozygous carriers of the rarer T-allele, lowest levels of oscillatory alpha power at most of the considered scalp sites were found. This observation was not verified by statistical testing because of the small number of subjects (also see Figure 6). Considering absolute alpha power density values, MANOVA did not demonstrate differences between genotype groups (F_(18,40)_ = 1.44, p = 0.165, Wilk’s Λ = 0.606).

**Figure 1.**
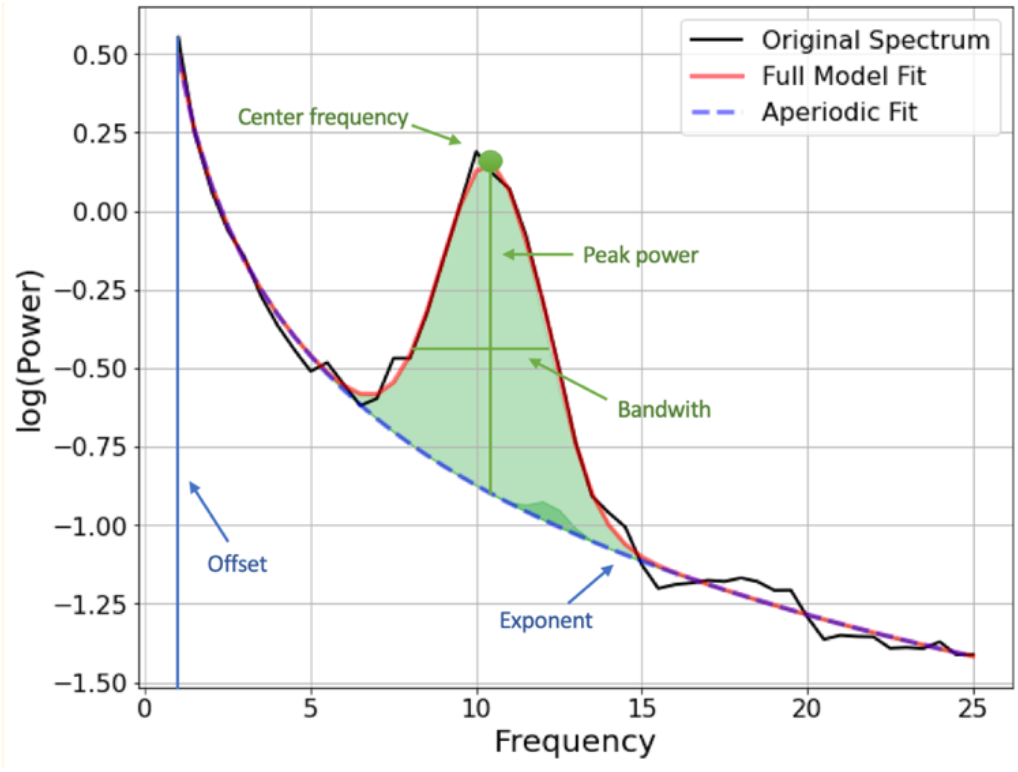
Individual power spectrum (at Oz) of an exemplary participant and FOOOF Fit

**Figure 2.**
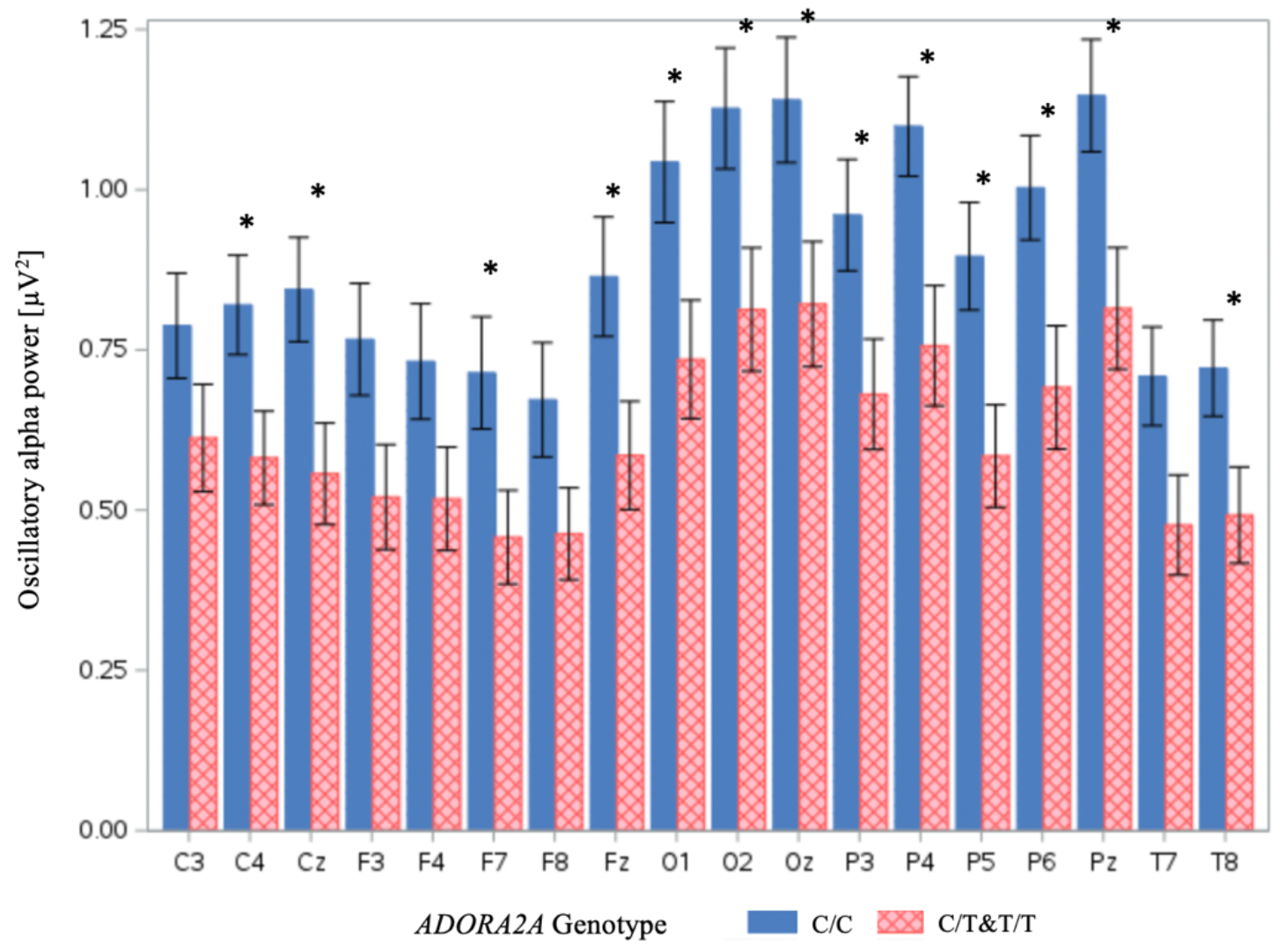
Waking EEG, mean oscillatory alpha power (and standard error) at different 10-20 system scalp derivations: Solid (blue)= C/C carriers of the ADORA2A polymorphism; Squared (red)= C/T and T/T carriers. Simple stars indicating significant difference after correction for multiple testing (p < 0.036, Benjamini Hochberg correction).

**Figure 3.**
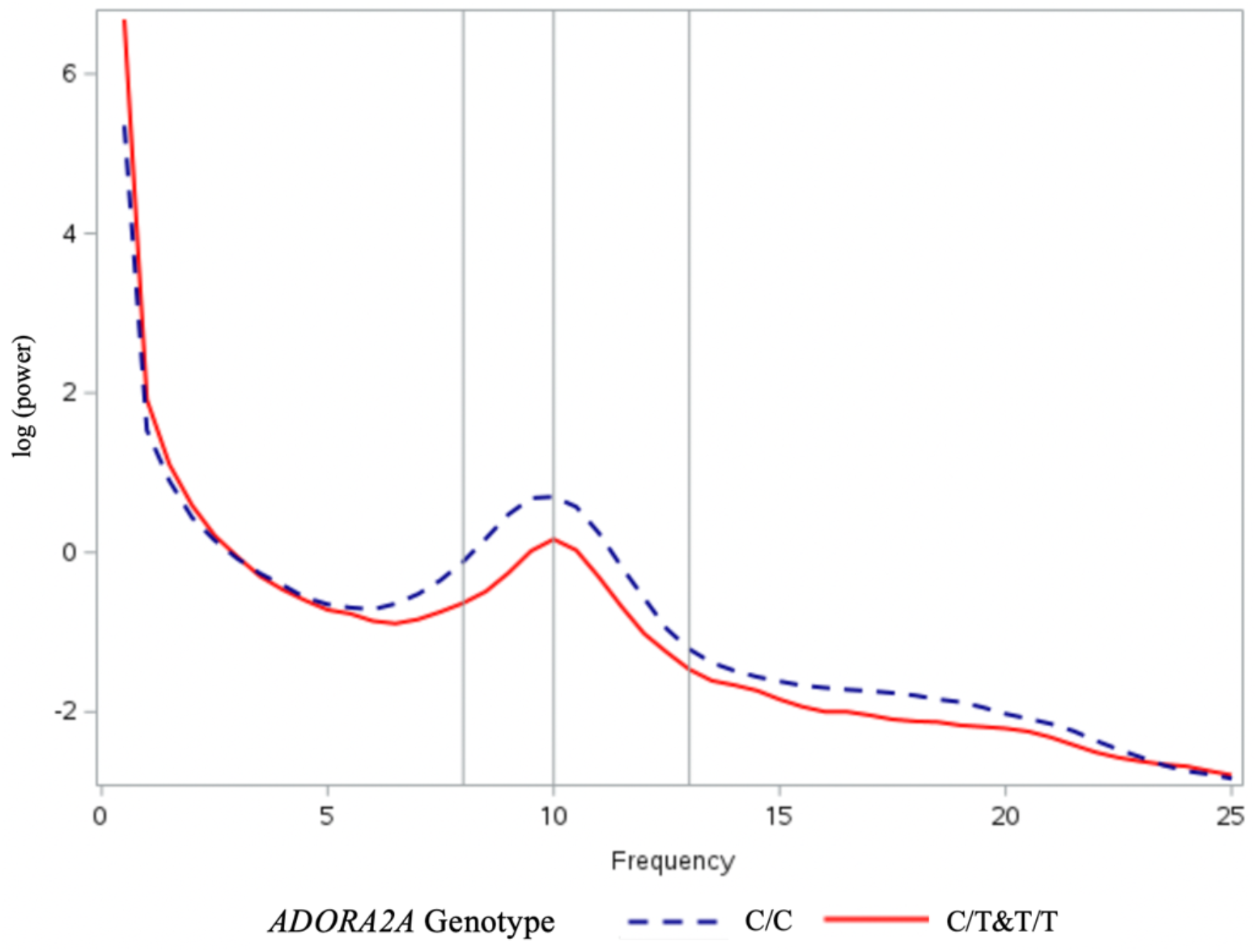
Wake EEG, geometric mean of power spectra at occipital derivations (Oz) Dashed (blue) = C/C carriers of the ADORA2A polymorphism; Solid (red) = C/T and T/T carriers.

**Figure 4.**
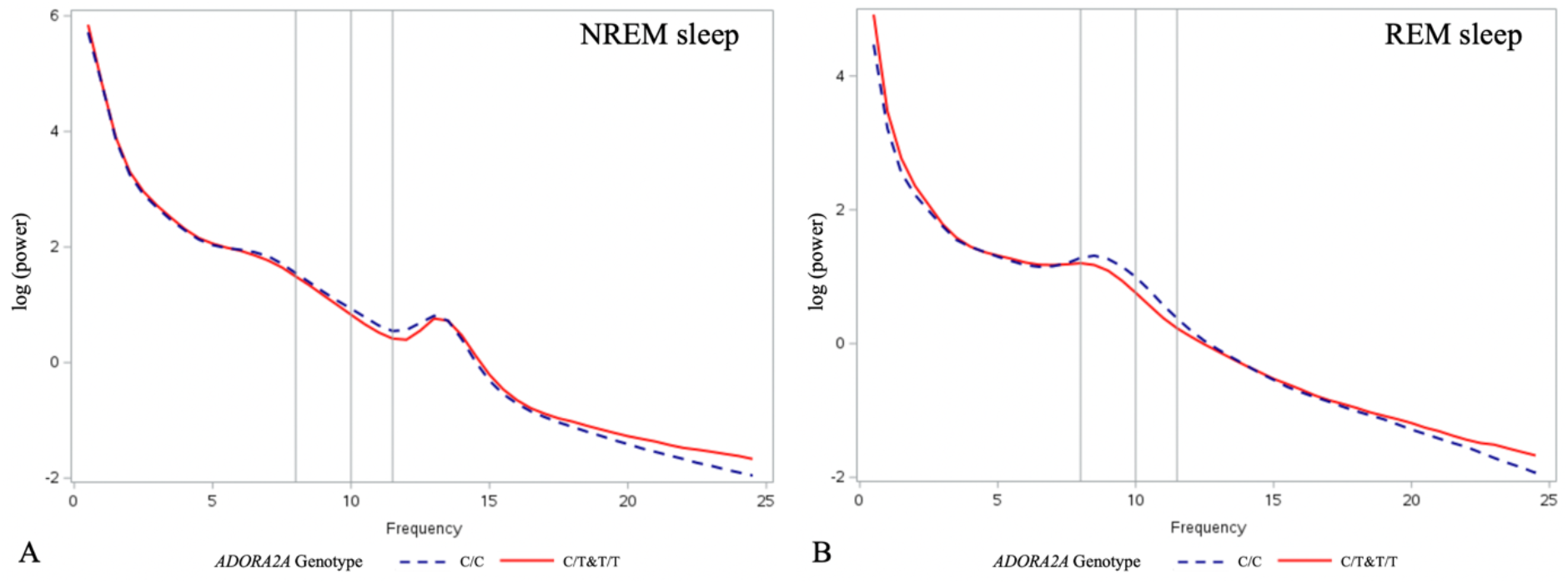
Sleep EEG (A NREM sleep, B REM sleep), geometric mean of absolute power spectra at occipital derivations (A1-O2, A2-O1): Dashed (blue) = C/C carriers of the ADORA2A polymorphism; Solid (red) = C/T and T/T carriers. Epochs of different sleep stages during sleeping period were cumulated.

**Figure 5.**
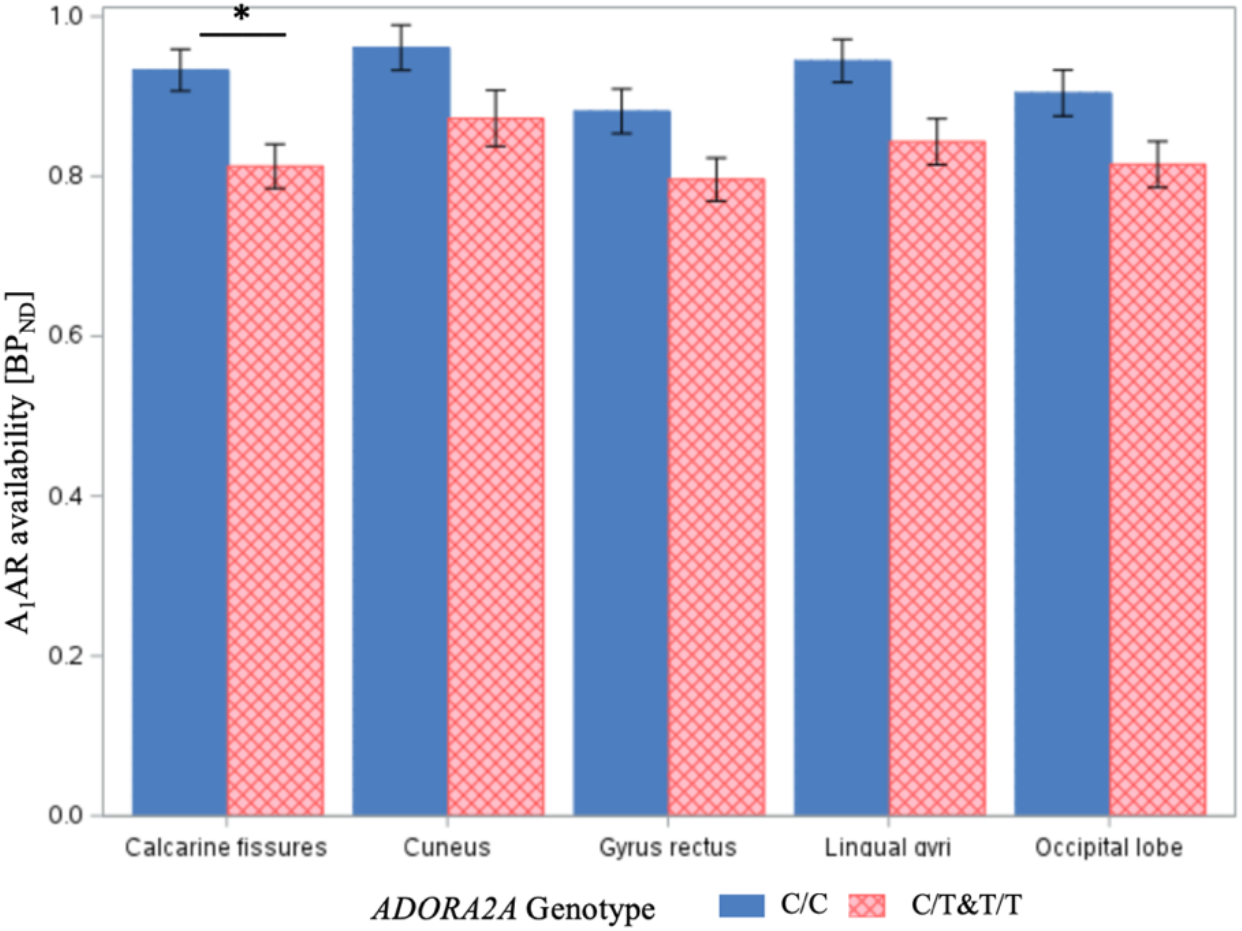
Mean A_1_AR binding potentials (and standard error) for genotype groups: Solid (blue) = C/C-allele carriers of the ADORA2A gene variant rs5751876 gene variant (n = 31); squared (red) = C/T-(n=23) and T/T-allele (n=5) carriers. In occipital brain regions C/C allele carriersindicate higher binding potentials compared to C/T- and T/T-carriers. Star indicates significant difference after correction for multiple testing (p <0.0025, Benjamini-Hochberg correction).

**Figure 6.**
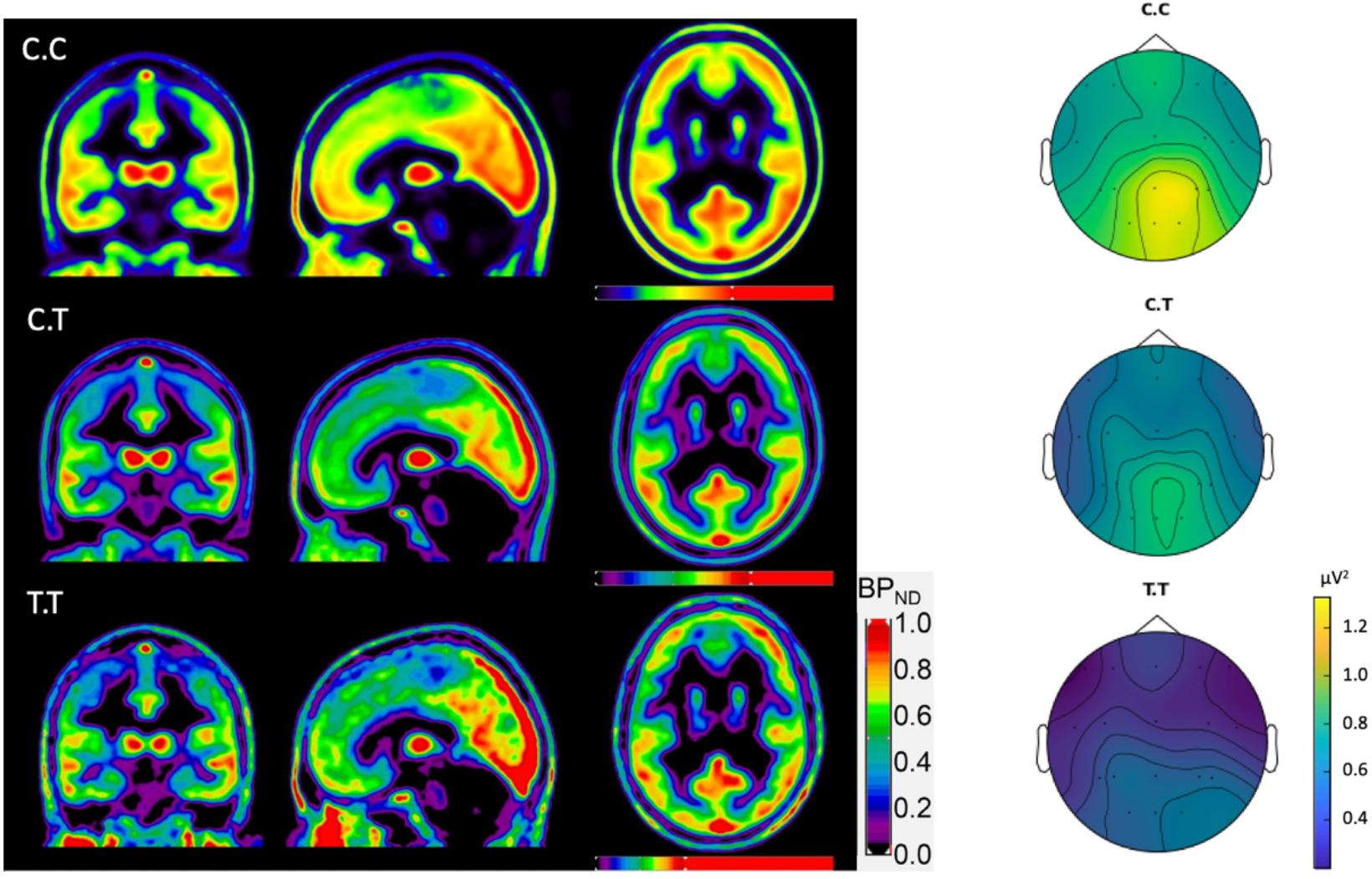
Mean A_1_AR binding potentials and averaged waking oscillatory alpha power (μV^2^) over the scalp in in C/C-(n = 29), C/T-(n = 22) and T/T-allele (n = 5) carriers of ADORA2A rs5751876 gene variant: The data of receptor availability and alpha power was averaged in each genotype separately. The T-allele seems to globally decrease alpha power derived over the scalp. The effect of genotype on receptor availability is less prominent.

The MANOVA of the sleep EEG data showed the difference in oscillatory alpha activity levels for different *ADORA2A* variant carriers to be consistent for REM sleep (overall effect F_(3,55)_ = 3.36, p = 0.025, Wilk’s Λ = 0.845), but not for NREM sleep (overall effect F_(3,55)_ = 0.89, p = 0.452, Wilks Λ= 0.954). Analysis of absolute alpha power density levels for sleep EEG, did not find differences between considered *ADORA2A* variant carriers (REM sleep: F_(3,55)_ = 2.16, p = 0.103, Wilk’s Λ = 0.895, NREM sleep: F_(3,55)_ = 0.47, p = 0.701).

Looking at the full power spectrum (see Figures 3 and 4) the alpha range between 8 and 13 Hz (8 and 11.5 Hz for sleep EEG) seems to be the frequency range where differences between genotype groups are most pronounced. Comparative analysis of other periodic alpha parameters like center frequency and bandwith as well as aperiodic parameters of offset and exponent did no indicate any significant differences between the considered groups

### 3.3 A_1_AR binding potentials in *ADORA2A* C/C- and T-allele carriers

The amount of A_1_AR receptors at rested state after a regular 8-hour sleep period was quantified. Highest BP_ND_ levels were found in occipital and temporal cortical regions as well as subcortical regions including thalamus. MANOVA demonstrated an overall difference between the two genotype groups (F_(18,40)_ = 2.62, p = 0.006, Wilk’s Λ = 0.459). In homozygous C/C-allele carriers (n = 31), slightly larger amounts of available A_1_AR receptors were quantified compared to C/T-(n = 23) and T/T-allele (n = 5) carriers (mean BP_ND_ in calcarine fissures 0.933 vs. 0.813). At uncorrected level this difference was seen in three occipital regions (calcarine fissures, lingual and occipital cortex), and the gyrus rectus (Figure 5). When considering single brain regions and correcting for multiple testing, however the effect was significant for the surrounding area of calcarine fissure only (p < 0.0025, Benjamini-Hochberg corrected significance level).

### 3.4 Regional oscillatory alpha powerduring waking and A_1_AR availability

Explorative correlation analysis was performed between waking oscillatory alpha power and A_1_AR BP_ND_. This included those brain regions and near-by electrodes, which showed most pronounced differences for different ADORA2A variant carriers. Also subcortical A_1_AR availability in the region of thalamus and oscillatory alpha power at central and occipital derivations were correlated. No associations between the amount of available A_1_AR in occipital brain region of calcarine fissure and oscillatory alpha power derived at Oz scalp derivation was found when including the complete sample (p = 0.078). Neither a correlation of A_1_AR binding potentials in subcortical region of thalamus and oscillatory alpha power at central scalp site (p = 0.075) or occipital scalp site (p = 0.365) was found. Because sex was previously demonstrated to modulate adenosine receptor availability (Hohoff et al., 2020; Pierling et al., 2021), we then analyzed the relationship of waking oscillatory alpha power and A_1_AR BP_ND_ in subgroups of females and males. The subgroup analysis revealed significant association between oscillatory alpha power at occipital scalp site and A_1_AR binding potential in occipital brain region of calcarine fissure in the sample of female participants (n = 24, r = 0.537, p = 0.007). However for the subgroup of males (n = 32, p = 0.409) no significant association were found (Figure 7). For occipital oscillatory alpha power extracted from epochs of wakefulness after sleep onset (polysomnography records) we found a similar correlation for female participants only (n = 26, r = 0.550, p = 0.004, A_1_AR availability in calcarine fissure).

**Figure 7.**
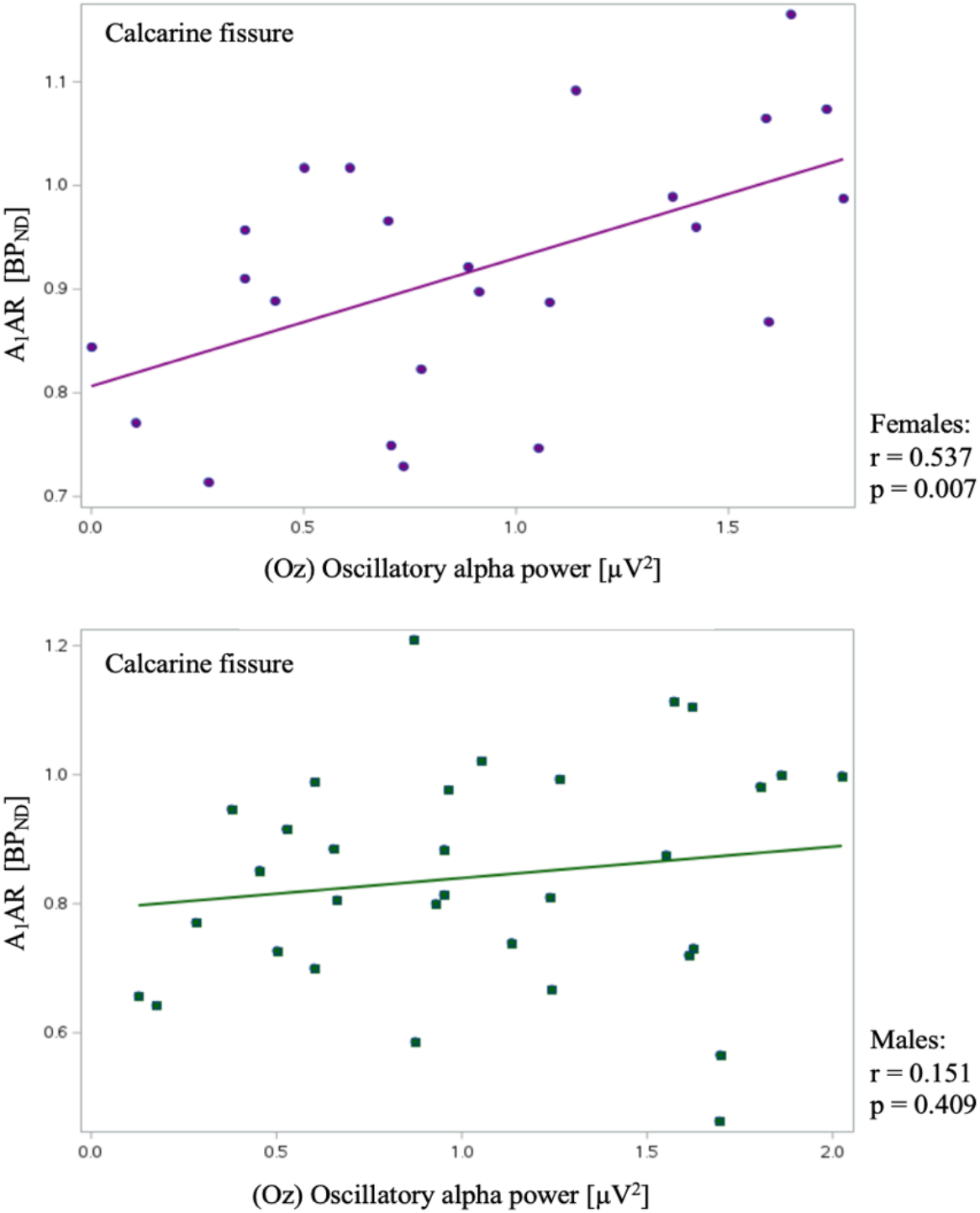
Subgroup analysis for correlation of A1AR availability in occipital brain region of calcarine fissure and oscillatory alpha power at occipital scalp site (Oz). Significant association for female participants (n = 24, r = 0.537, p= 0.007), but not for male participants (n = 32, r = 0.151, p = 0.409)

## 4. Discussion

In this study EEG alpha activity in the state of rested wakefulness and during an 8-hour sleep period as well as cerebral A_1_AR availability were assessed during a controlled laboratory setting for the first time. We aimed to investigate possible influences of adenosine signaling via A_1_AR on alpha power derived at scalp level. As genetic variation of the gene encoding the adenosine receptor type 2A (*ADORA2A*) was shown to modulate not only A_1_AR availability but also EEG correlates, we explored differences in these two parameters in a large sample of healthy, right-handed, genotyped subjects.

We found homozygous C-allele carriers of the *ADORA2A* gene variant to indicate higher oscillatory alpha power values than homo- and heterozygous carriers of the T-allele during wake state as well as in REM sleep. These results correspond with higher alpha power found by Rétey and colleagues in C/C-allele carriers compared to T/T - allele carriers, analyzing absolute EEG power derived at one bipolar C3-A2 derivation. Differences in EEG alpha power were demonstrated for wake state and during REM as well as during NREM sleep (Retey et al., 2005). The present analysis shows that this is a global effect, as differences were found for 13 of the 18 considered scalp sites. As homozygous T-allele carriers indicated lowest and homozygote C-allele carriers highest alpha power density levels, a gene-dose relation can be assumed. EEG variables such as alpha power were shown to be a rather stable individual trait, which is genetically inherited (van Beijsterveldt & van Baal, 2002). Thus, a modulatory effect of *ADORA2A* could be a component in a pattern of genetic variance that forms the basis for these electroencephalographic phenotypes. We did not find higher oscillatory alpha power values in C/C-allele carriers during NREM sleep. This may reflect state-dependant power changes in different frequency ranges and separate underlying generation processes. Measured mean oscillatory alpha power decreased generally with increasing sleep depth and differences between genotype-groups diminished. During NREM sleep, cerebral blood flow is reduced, and this was shown to accompany alterations in neuronal transmission and excitability (Bazanova & Vernon, 2014; Kotajima et al., 2005). Interestingly we could demonstrate differences in alpha activity between *ADORA2A* variant carriers only for oscillatory alpha power extracted using FOOOF tool, not absolute power density estimates. Acoordingly, separating aperiodic and periodic component of neural power spectra may help to differentiate possible effects of genetical influence on EEG parameters.

A modulatory effect of *ADORA2A* polymorphism on cerebral A_1_AR availability was confirmed, although the differences in adenosine receptor availability between gene allele carriers was smaller than previously suggested by Hohoff and colleagues (Hohoff et al., 2020). The authors discussed several confounding factors like sex, prior sleep amount, coffee intake and smoking behavior. Additionally, influence of other gene polymorphisms of the adenosinergic and dopaminergic transmitter systems were investigated. In the present study, sex and age were equally distributed between the two compared groups. During sleep, adenosine concentration decreases and concurrently A_1_AR availability is returned to baseline levels (Elmenhorst et al., 2017). Sleep duration in the night before the measurement was scheduled and assessed with polysomnography to minimize differences between subjects. Nonetheless, inter-individual preferences and habits regarding sleep amount were not considered, which means that differences in individual A_1_AR availability might be masked or blurred as some subjects did sleep longer or shorter than usual.

Waking alpha activity with its inhibitory function modulates neuronal excitability (Romei et al., 2008) and is interrelated to cellular metabolism. Variation in the amount of measured alpha power may be due to phasic changes, i.e., rapid and direct modulation by attentional processes or tonic changes embracing arousal processes like homeostatic and circadian sleep drive, aging and neurodegenerative processes (Klimesch, 1999). In this analysis, an average of two records on the same day, each lasting several minutes, was considered to compensate for transient fluctuations related to changes in attentiveness. Considering the whole sample we did not find associations between binding potentials of A_1_AR and derived waking oscillatory alpha power density at nearby scalp sites. Neither a quantitative relation between A_1_AR availability in the thalamus and derived oscillatory alpha power at the scalp was found. A correlation between occipital oscillatory alpha power and A_1_AR availability was demonstrated only for the subgroup of females (n = 24). This might indicate differential alpha modulation for females and males. Indeed there is a growing knowledge on sex differences in brain organization, including brain structure, function and neurotransmitter-systems of serotonin, dopamine and other (Cosgrove et al., 2007). In addition, gonadal hormones are known to affect neural functioning (McEwen & Milner, 2017). As we did not collect any information about the phase of menstrual cycle of the participants, this limits the investigation of hormonal influences. Several studies investigating alpha activity in relation to the blood oxygen level dependent (BOLD) signal with functional MR-imaging showed modulations in EEG alpha rhythm to be correlated with BOLD changes in the cortex (Goncalves et al., 2006; Knaut et al., 2019; Laufs et al., 2003) and the thalamus (Goldman et al., 2002; Moosmann et al., 2003; Sadaghiani et al., 2010). Studies applying [^18^F]Fluordesoxyglucose (FDG) PET found alpha power to be correlated with glucose metabolism in thalamic regions (Larson et al., 1998; Lindgren et al., 1999). These findings suggest that ‘activation’, represented by increase of oxygen usage (in fMRI) or glucose depletion (in FDG-PET), of the cortex is followed by alpha suppression and vice versa. For thalamic regions, both positive and negative correlations were found (Moosmann et al., 2003), suggesting a more complex role in alpha generation. However, a simple bi-directional relation cannot be assumed, as pharmacological studies investigating agents such as benzodiazepines that suppress neuronal metabolism have found a concomitant decrease in alpha activity (Lozano-Soldevilla, 2018).

To fight fatigue, caffeine is a very popular and commonly consumed substance. Its wake-promoting effect is mediated via unselective blocking of adenosine receptors. Coffee administration was shown to be followed by a general decrease in alpha power (Barry et al., 2005). Adenosine accumulates as a result of several metabolic pathways and is catabolized among others by the enzyme adenosine deaminase (ADA) (Urry & Landolt, 2015). It was demonstrated that individuals with genetic variation of ADA, resulting in higher endogenous concentrations of the transmitter adenosine, have higher levels of slow wave activity during sleep (Retey et al., 2005). Adenosine, as a homeostatic sleep factor, influences arousal status and cellular effects including a decrease of neuronal excitability (Dulla et al., 2005). In fact, different functions of A_1_ARs and A_2A_ARs in sleep-wake regulation are discussed. While sleep onset might be initiated via A_2A_AR transmission, adenosine signaling via A_1_ARs is suggested to maintain sleep state. Wake promoting neuron groups located in the basal forebrain are inhibited via A_1_AR stimulation, whereas sleep promoting neurons are disinhibited (Lazarus et al., 2019). We assume global effects of adenosine signaling on EEG alpha power to act via the A_1_AR as this receptor type is ubiquitously distributed in the cortex unlike the A_2_AR, which is mostly concentrated in subcortical regions including striatum and hypothalamus (Huang et al., 2011). Adenosine was shown to have an inhibititory tone on mesopontine cholinergic neurons, which increase EEG patterns of arousal (Rainnie et al., 1994). Next to that adenosine inhibits via A_1_AR cortical acetylcholine release (Materi et al., 2000). This might be pathways by which the receptor availability contributes to individual differences in oscillatory alpha power at occipital scalp site. Yet further investigation is needed, especially to elucidate the suggested sex differences in adenosinergic effects on individual EEG alpha activity.

## 5. Conclusion

EEG parameters like oscillatory alpha power present intra-individual stability and inter-individual variation. Here, we showed that variation of the *ADORA2A* gene is related to inter-individual variation of oscillatory alpha power recorded during rested wakefulness as well as during REM sleep. Furthermore, *ADORA2A* gene variation modulates the availability of cerebral A_1_ARs. The subgroup of female participants indicated an association between occipital A_1_AR availability and occipital oscillatory alpha power. In summary, individual alpha formation seems to be subject to genetic influence, including variations in the adenosine system, although evidence for a direct modulation via A_1_AR was found only for females.

## 6. Declarations

### 6.1 Data and code availability statement

All MR and PET and EEG data analysed during the current studies are available from the corresponding author on reasonable request.

### 6.4 Ethics approval

All procedures performed in studies involving human participants were in accordance with ethical standards of the institutional and/or national research committee and with the 1964 Helsinki declaration and its later amendments or comparable ethical standards. The studies were approved by the Ethics Committee of the regional Medical Board (Ärztekammer Nordrein) and the German Federal Office for Radiation Protection.

### 6.5 Consent of participate

Informed consent was obtained from all individual participants included in the studies.

### 6.6 German Clinical Trial Registry

DRKS #DRKS00010194, registered 22 March 2016, https://www.drks.de/drks_web/navigate.do?navigationId=trial.HTML&TRIAL_ID=DRKS00010194 DRKS #DRKS00014379, registered 04 April 2018, https://www.drks.de/drks_web/navigate.do?navigationId=trial.HTML&TRIAL_ID=DRKS00014379

## 7. Acknowledgements

We thank all volunteers for participating in the studies, Sylvia Köhler-Dibowski and Stephanie Krause from the Forschungszentrum Jülich and Annette von Waechter of the German Aerospace Center for their excellent technical assistance and support in study conductance.

